# High-risk human papillomaviruses down-regulate expression of the Ste20 family kinase MST1 to inhibit the Hippo pathway and promote transformation

**DOI:** 10.1101/369447

**Authors:** Ethan L. Morgan, Molly R. Patterson, Siu Yi Lee, Christopher W. Wasson, Andrew Macdonald

## Abstract

Human papillomaviruses (HPV) are a major cause of malignancy worldwide They are the aetiological agent of almost all cervical cancers and an increasing number of head and neck carcinomas. Deregulation of the Hippo pathway component YAP1 has recently been demonstrated to play a role in HPV-mediated cervical cancer, but whether other components of this pathway are implicated in the pathogenesis of this disease remains poorly understood.

The expression level and activation status of critical Hippo pathway components were analysed across multiple cytology samples from patients with cervical disease, as well as HPV positive (HPV+) and HPV negative (HPV-) cervical cancer cell lines using real time qPCR, western blot and immunohistochemistry. In parallel, we assessed the effects of MST1 and MST2 overexpression upon cervical cancer cell proliferation, migration and invasion. Finally, we interrogated the consequences of interrupted MST1 and MST2 function using a targeted small molecule inhibitor in tandem with kinase inactive MST mutants. Our analysis found that expression of the Ste20 kinase MST1 was decreased within both HPV+ primary patient samples and cervical cancer cell lines. This effect was mediated by the virus-coded oncoproteins E6 and E7, which impair MST1 transcription. Reintroduction of MST1, or its paralogue MST2, into HPV positive cervical cancer cells re-activated the Hippo pathway, leading to a reduction in cell proliferation, migration and invasion. Finally, using a small molecule inhibitor of MST1/2 or kinase inactive mutants of either protein, we demonstrated that this effect required the kinase function of MST1/2. Our results reveal that HPV down regulates MST1 expression to inactivate the Hippo pathway and so drive cells towards transformation.

## Introduction

Human papillomaviruses (HPV) are the major cause of cervical cancer, accounting for 99.9% of all cases, as well as being linked to an increasing number of head and neck carcinomas (1-3). The majority of HPV-associated cancer cases are caused by infections with two high-risk (HR) types; HPV16 and HPV18, but other HPV types can also cause cancer. The main drivers of transformation in HPV-mediated cancers are the E6 and E7 oncoproteins. These oncoproteins interact with multiple host cell factors to manipulate cellular processes and promote transformation (4-7).

The Hippo pathway is a major regulator of cellular homeostasis. In humans, the canonical Hippo pathway consists of the Ste20 family kinases MST1 and MST2, which are activated by phosphorylation in response to a wide range of upstream factors (8). Activated MST1/2 subsequently phosphorylate the adaptor protein MOB1 and the Ste20 kinases LATS1 and LATS2, which in turn phosphorylate the transcriptional regulators YAP1 and TAZ (9). Phosphorylation of YAP1 results in its nuclear exclusion or proteosomal degradation (9). However, in their non-phosphorylated state, YAP1/TAZ traffic to the nucleus where they partner with TEAD transcription factors to regulate the expression of genes controlling cell proliferation (10).

The *YAP* gene is a *bona fide* oncogene, with a number of studies showing its importance as a growth promoter (11,12). *YAP1* is overexpressed in several cancers, with increased expression linked to shorter patient survival (13-15). Studies also show that loss of MST1/2 contributes to tumourigenesis. In mice, reduced MST1/2 mRNA levels lead to hepatocellular carcinoma (16) and in colorectal cancers, expression of micro-RNA miR-590-3p decreases MST1 expression (17). One recent study recommended MST1 as an early biomarker for colorectal cancer and as a marker for poor prognosis (18).

Despite the emerging paradigm of the Hippo pathway as a key mediator of transformation, its role in cervical cancer is poorly understood. Recently, it was demonstrated that the HPV E6 oncoprotein stabilises YAP protein expression, thereby promoting cervical cancer progression (19). YAP stabilisation leads to an increased expression of the epidermal growth factor receptor (EGFR) ligand, amphiregulin (AREG), which in turn further inhibits the Hippo pathway (19). Whether other components of the Hippo pathway are deregulated in cervical cancer is less clear.

To gain further insights into the regulation of the Hippo pathway in cervical cancer we firstly analysed the expression and activation of key components of the pathway in a panel of cervical cancer cell lines. We demonstrated that MST1 expression is selectively decreased in HPV positive (HPV+) cervical cancer cells when compared with normal human keratinocytes (NHKs) or HPV negative (HPV-) cervical cancer cells. Importantly, we showed that MST1 expression is also decreased in cytology samples from patients with cervical disease and in cervical cancer samples. Additionally, reintroduction of MST1, or its paralogue MST2, into HPV+ cancer cells inhibits their proliferation, migration and invasion. Finally, we confirm that the tumour suppressor activity of MST1 is dependent on its function as a protein kinase. Together, we have identified a novel host factor targeted by HPV to drive cervical cancer progression that may be a potential therapeutic target.

## Materials and Methods

### Cell culture

HeLa (HPV18+ cervical epithelial adenocarcinoma cells), SW756 (HPV18+ cervical squamous carcinoma cells), SiHa (HPV16+ cervical squamous carcinoma cells), CaSKi (HPV16+ cervical squamous carcinoma cells) and C33A (HPV-cervical squamous carcinoma) cells obtained from the ATCC were grown in DMEM supplemented with 10% FBS (ThermoFischer Scientific, USA) and 50 U/mL penicillin. Normal human keratinocytes (NHKs) were maintained in serum free medium (SFM; GIBCO, UK) supplemented with 25 μg/mL bovine pituitary extract (GIBCO, UK) and 0.2 ng/mL recombinant EGF (GIBCO, UK). All cells were cultured at 37°C and 5% CO_2_. Cells were negative for Mycoplasma contamination during this investigation. Cell identify was confirmed by karyotype analysis.

### Plasmids and inhibitors

pIC-MST1-Myc and pIC-MST2-Myc were provided by Steve Cohen, LMCB, Singapore. pJ3M-MST1 K59R (Addgene plasmid # 12204) and pJ3H-MST2 K56R (Addgene plasmid # 12206) were purchased from Addgene (Cambridge, MA, USA). The small molecule inhibitor XMU-MP1 was used at a final dose of 1μM (20).

### Transfections and mammalian cell lysis

Transfections were performed with a DNA to Lipofectamine^®^ 2000 (ThermoFischer) ratio of 1:2.5. 48 hr post transfection, cells were lysed in lysis buffer for western blot analysis (6).

### Western blot analysis

Equal amounts of protein from cell lysates were separated by SDS PAGE and transferred onto a nitrocellulose membrane by a semi-dry transfer method (Trans Blot^®^ SD Semi-Dry Transfer cell, Bio-Rad, USA). Membranes were blocked with 5 % milk solution before incubation with primary antibodies at 1:1000 dilution unless otherwise stated: p-YAP1 Ser-127 (CST, D9W2I), YAP1 (CST, D8H1X), p-MOB1 (CST D2F1O), MOB1 (CST E1N9D), MST1 (Abcam ab51134) (1:2000), MST2 (Abcam, ab52641) (1:5000), Myc (9E10), Cyclin B1 (Santa Cruz Biotechnology (SCBT); sc-245), Cyclin D1 (Abcam ab13475), Cyclin E (Thermofisher, 32-1600), HPV 16/18 E6 (CBT; sc-460), HPV 16 E7 (SCBT; sc-1587), HPV 18 E7 (Abcam ab100953), and GAPDH (SCBT; sc365062) (1:5000) as a loading control. Horseradish peroxidase (HRP)-conjugated secondary antibodies (Sigma, USA) were used at a 1:5000 dilution. Proteins were detected using WesternBright ECL (Advansta, USA) and visualised on X-ray film.

### RNA extraction, cDNA synthesis and quantitative Real Time-PCR

Total RNA was extracted from cells using an E.Z.N.A Total RNA Kit I (omegabiotek). cDNA was synthesised using 1 μg of RNA and iScript cDNA synthesis kit (Bio Rad). qRT-PCR was performed on the synthesised cDNA on a Corbett Rotor-Gene 6000 using QuantiFast SYBR Green PCR kit (Qiagen) and analysed using the ∆∆CT method(21) normalised to the U6 housekeeping gene. Each experiment was repeated at least 3 times. Primer sequences can be found in Table 1.

**Table 1.**
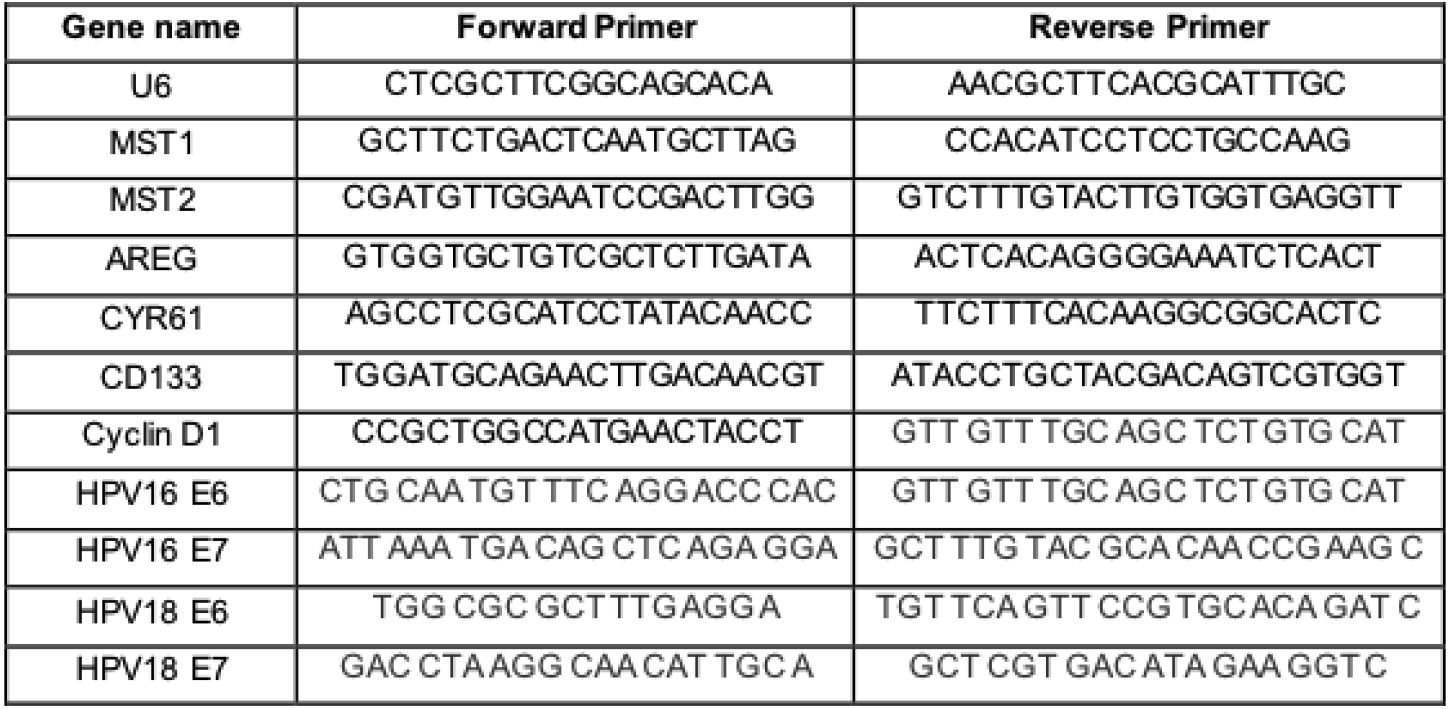

### Colony formation assay

48 hr post-transfection, cells were trypsinised and reseeded in a six well plate at 500 cells per well and left to incubate for 14 - 21 days. Colonies were then stained (1 % crystal violet, 25 % methanol) and colonies were counted manually. Each experiment was repeated a minimum of 3 times.

### Soft agar assay

Cells were transfected as required. 60 mm dishes were coated with a layer of 1 % agarose (ThermoFischer Scientific, USA) in 2X DMEM (ThermoFischer Scientific, USA) supplemented with 20 % FBS. 48 hr post-transfection, cells were trypsinised and added to 0.7 % agarose in 2X DMEM (ThermoFischer Scientific, USA) supplemented with 20 % FBS at 1000 cells/mL. Once set, DMEM supplemented with 10 % FBS and 50 U/mL penicillin was added. The plates were then incubated for 14 - 21 days. Each experiment was repeated at least three times unless stated otherwise. Visible colonies were counted manually.

### Flow cytometry

Cells were transfected as required. 48 hr post-transfection, cells were harvested and fixed in 70 % ethanol overnight. The ethanol was removed and cells washed with PBS containing 0.5 % (w/v) BSA. Cells were stained with PBS containing 0.5 % BSA, 50 μg/mL propidium iodide (Sigma) and 5 μg/mL RNase (Sigma) and incubated in this solution for 30 min at room temperature. Samples were processed on an LSRFortessaTM cell analyzer (BD) and the PI histograms analysed on modifit software. Each experiment was repeated a minimum of 3 times.

### ELISA

Human amphiregulin levels were detected in the cell supernatants using the DuoSet^®^ ELISA kit according to the manufacturer’s instructions (R&D Systems).

### Microarray analysis

For microarray analysis, a dataset of 28 cervical cancer cases and 23 normal cervix samples was utilised. Microarray data was obtained from GEO database accession number GSE9750.

### Immunofluorescence Analysis

Cells were seeded onto coverslips and, 24 hr later, were transfected as required. 24 hr after transfection, cells were fixed with 4 % paraformaldehyde for 10 min and then permeabilised with 0.1 % (v/v) Triton for 15 minutes. Cells were then incubated in primary antibodies in PBS with 4 % BSA overnight at 4°C. Primary antibodies were used at a concentration of 1:400. Cells were washed thoroughly in PBS and then incubated with Alex-fluor conjugated secondary antibodies 594 and Alexa 488 (1:1000) (Invitrogen) in PBS with 4% BSA for 2 hours. DAPI was used to visualise nuclei. Coverslips were mounted onto slides with Prolong Gold (Invitrogen).

### Transwell^®^ migration and Invasion assays

Cells were transfected as required. 48 hr post-transfection, cells were harvested and resuspended in serum free DMEM. 5×10^4^ cells were replated onto the upper chamber of a Transwell^®^ filter with 8 μm pores (Falcon^®^, Corning, USA) coated with (invasion) or without (migration) 200 μg/mL Matrigel^®^ Matrix (Corning). DMEM containing 10% FBS was used as the chemoattractant. After 24 hr, non-migrated cells on the upper side of the filter were removed with a cotton swab, and cells on the underside of the filter were stained with 1% crystal violet in 25% methanol. For each experiment, the number of cells in five random fields on the underside of the filter was counted, and three independent filters were analysed.

### Statistical Analysis

Where indicated, data was analysed using a two-tailed, unpaired Student’s t-test.

## Results

### MST1 expression is selectively decreased in HPV+ cervical cancer cell lines

To investigate the regulation of the Hippo pathway in cervical cancer cells, we first measured the protein expression and enzymatic activity of key components of the pathway. Compared with normal primary human keratinocytes (NHKs) or the HPV-cervical cancer cell line C33A, levels of total YAP1 and the phosphorylated form of YAP1 (pYAP1 S127) were significantly increased (Figure 1A), consistent with a previous study linking increased YAP1 expression to cervical cancer(22). In these experiments we also observed that MST1 protein expression was significantly lower in both HPV16+ (SiHa p= 0.0087 and Caski p= 0.0044) and HPV18+ (SW756 p= 0.0052 and HeLa p= 0.0018) cervical cancer cell lines compared to NHK cells. Levels of MST1 in the C33A cells were similar to those observed in the NHK cells (p= 0.44) (Figure 1A, quantified in B). The reduction in MST1 protein expression in the HPV+ cell lines was accompanied by a decrease in phosphorylation of the MST1 substrate MOB1 (Figure 1A).

**Figure 1.**
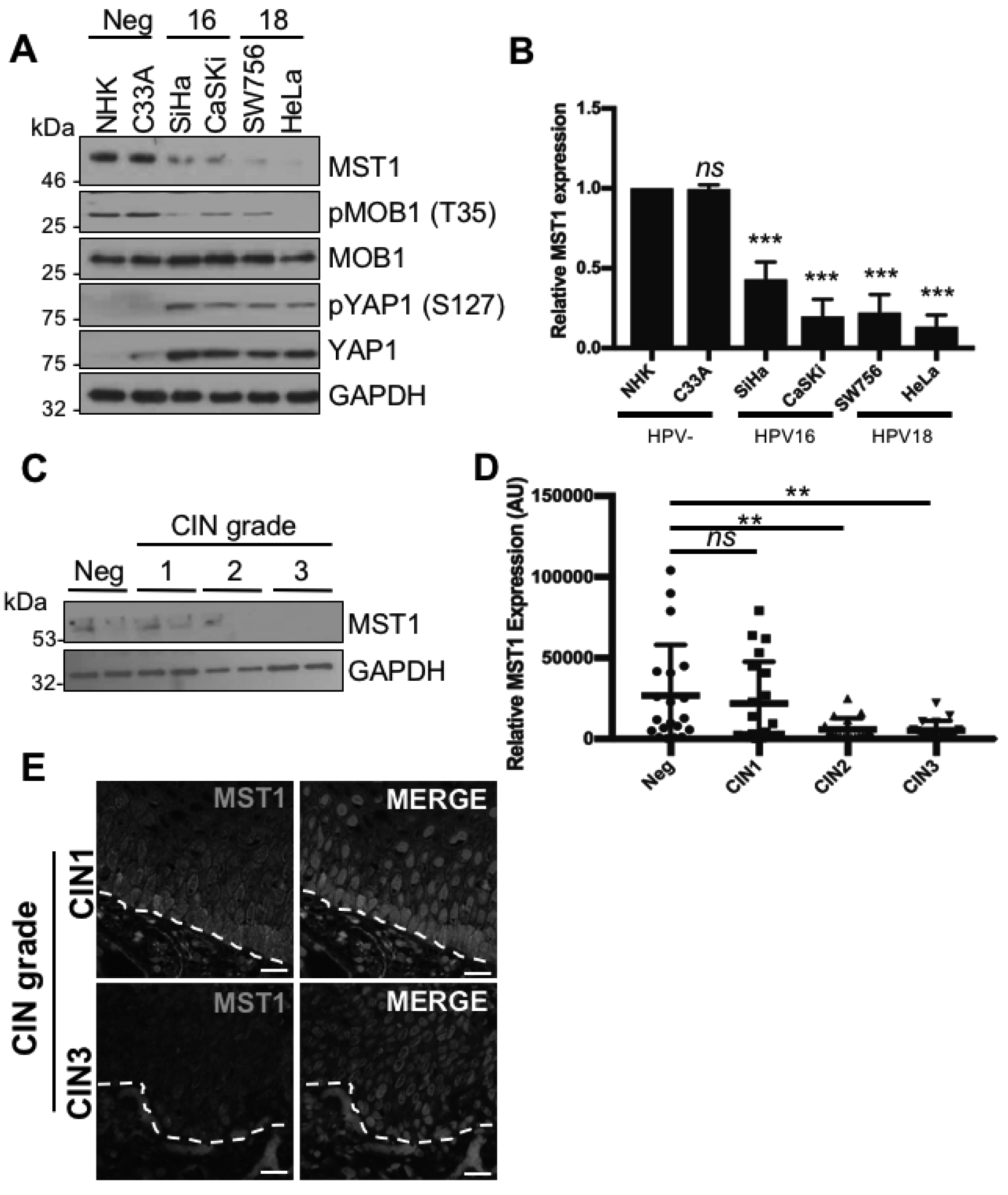
MST1 expression is down regulated in HPV+ cervical cancer. **A)** Representative western blot analysis of a panel of HPV- and HPV+ cervical cancer cell lines for components of the Hippo pathway. GAPDH served as a loading control. (N=3). **B)** Densitometry analysis of MST1 expression in **A** from four independent experiments. **C)** Representative western blot from cytology samples of CIN lesions of increasing grade analysed for MST1 expression. GAPDH served as a loading control. **D)** Scatter plot of densitometry analysis of a panel of cytology samples. Twenty samples from each clinical grade (Neg, CIN 1-3) were analysed by western blot and densitometry analysed was performed for MST1 expression. **E)** Representative immunofluorescence analysis of tissue sections from cervical lesions representing LSIL and HSIL. Sections were stained for MST1 expression (green) and nuclei were visualised using DAPI (blue). Images were acquired using identical exposure times. Scale bar, 20 μm. Error bars represent the mean +/- standard deviation of a minimum of three biological repeats. *P<0.05, **P<0.01, ***P<0.001 (Student’s t-test).

To extend these results to a physiologically relevant scenario we analysed cervical liquid based cytology samples from a cohort of HPV16+ patients representing the progression of disease development. Cervical intraepithelial neoplasia (CIN) commonly precedes cervical cancer progression. CIN1 is thought to represent a transient HPV infection, while CIN3 represents clinically significant HPV infection that may progress to cervical cancer (23). Normal cytology samples from HPV-patients served as controls in these experiments. By western blot we observed a significant decrease in MST1 protein levels during cervical disease progression through the CIN grades (Figure 1C and quantified in 1D; CIN1 p>0.05; CIN2 p=0.012; CIN3 p=0.015). We corroborated these findings using immunofluorescence microscopy to examine MST1 protein expression in sections of cervical tissue from both low-grade CIN1 and high-grade CIN3 samples. Analysis of the staining pattern showed a clear decrease in MST1 protein staining in the CIN3 section; in agreement with the western blot data (Figure 1E). Together, these data demonstrate that MST1 expression is selectively reduced in HPV+ disease.

### MST1 expression is inhibited at the transcript level in HPV+ cervical cancer cells

To investigate the mechanism by which MST1 expression is decreased in HPV+ cervical cancer cells, we measured MST1 mRNA levels using qRT-PCR (Figure 2A). MST1 mRNA levels were selectively decreased in both HPV16+ (SiHa p= 0.01 and CaSKi p= 0.004) and HPV18+ (SW756 p= 0.001 and HeLa p= 0.05) cervical cancer cell lines when compared with NHK and C33A cells (p= 0.12). Next, we extracted RNA from cervical cytology samples and performed qRT-PCR analysis to determine the levels of MST1 mRNA. In agreement with the protein data, there was a significant decrease in MST1 mRNA levels during cervical disease progression (Figure 2B; CIN1 p>0.05; CIN2 p=4.6×10^-7^; CIN3 p=1.1×10^-5^). Finally, we mined a publically available database of microarray data comparing normal cervical samples against those obtained from cervical cancer samples(24). In these samples, there was a statistically significant decrease in MST1 mRNA expression in the cervical cancer samples (Figure 2C; p=0.01). Together, these data demonstrated that MST1 levels are regulated at the mRNA level in HPV+ cervical cancer.

**Figure 2.**
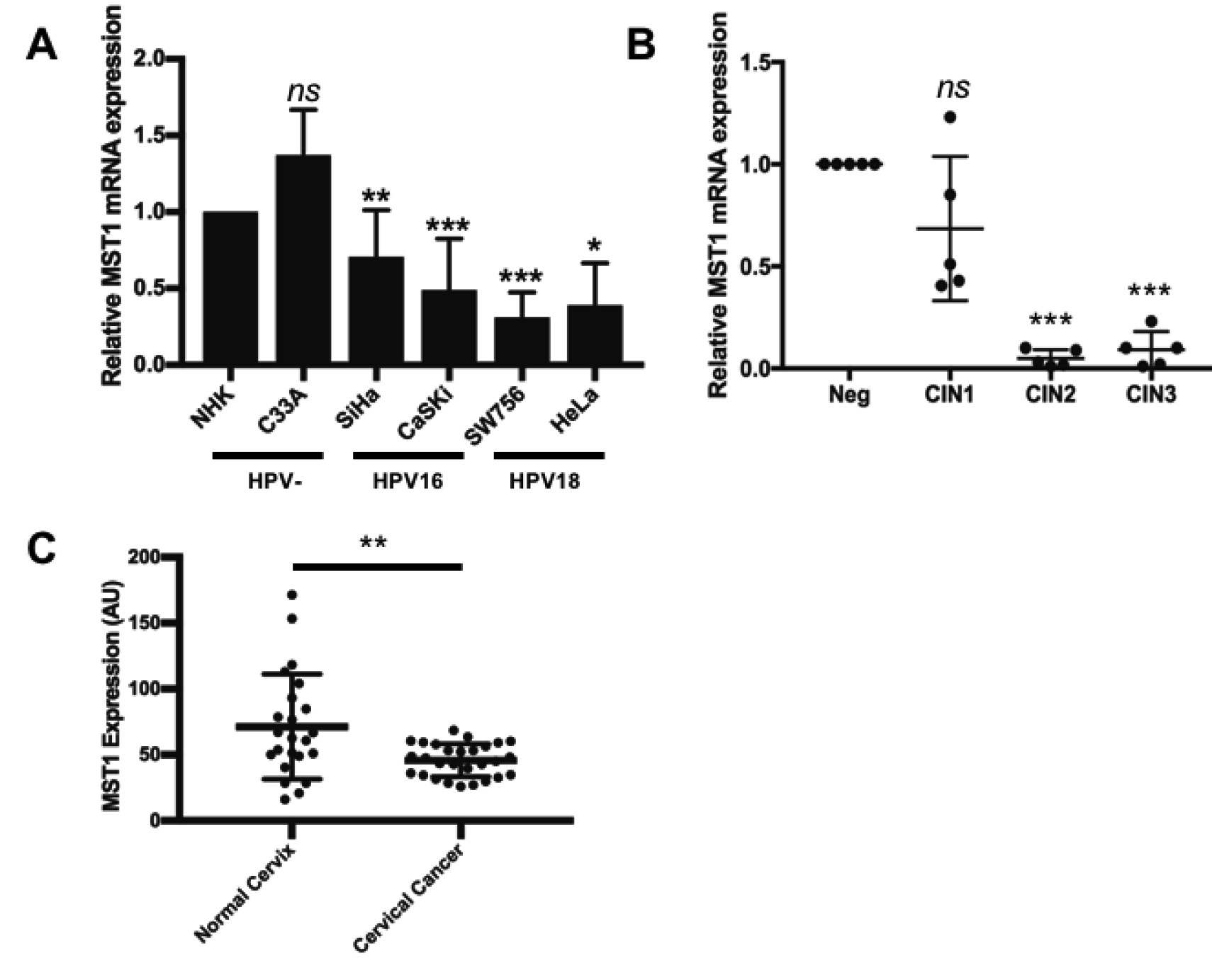
MST1 is regulated at the transcript level in HPV+ cervical cancer. **A)** qPCR analysis of MST1 transcript levels in a panel of HPV- and HPV+ cervical cancer cell lines. U6 expression was used as a loading control (N=3). **B)** qPCR analysis of MST1 transcript levels from different CIN grades (n=5 from each grade). U6 expression was used as a loading control. **C)** Scatter dot plot of data acquired from the dataset GSE9750 on the GEO database. Arbitrary values for the mRNA expression of MST1 in normal cervix (n=23) and cervical cancer (n=28) samples were plotted. Error bars represent the mean +/- standard deviation of a minimum of three biological repeats. *P<0.05, **P<0.01, ***P<0.001 (Student’s t-test).

### HPV E6 and E7 oncoproteins inhibit MST1 expression

The E6 and E7 oncoproteins are the key drivers of transformation in HPV+ cervical cancers(6,25,26). We hypothesized that one of these oncoproteins may be responsible for the reduced MST1 expression observed in cancer cell lines and clinical samples. To investigate this, we employed a siRNA knockdown strategy in which we transfected HPV18+ HeLa or HPV16+ CaSKi cells with a pool of specific siRNAs targeting E6 and E7 alone or in combination(6,27). Transfection of siRNA targeting E6 resulted in the loss of E6 protein expression and a small reduction in E7 protein levels in both cell lines (Figure 3A; lane 3). E6 knockdown resulted in the re-appearance of MST1 protein expression (Figure 3A; lane 3 and quantification in Figure 3B) in both cancer cell lines. SiRNA mediated knockdown of HPV E7 also increased MST1 protein expression, although the effect was less pronounced than for E6 knockdown (Figure 3A; lane 4 and quantification in Figure 3B). The combined knockdown of both E6 and E7 led to the most substantial increase in MST1 expression (Figure 3A; lane 5 and quantification in Figure 3B). We next used qRT-PCR to confirm if E6 and E7 regulate MST1 mRNA expression. In agreement with our previous data, combination of E6 and E7 knockdown led to a significant increase in MST1 mRNA expression in both HPV16+ and HPV18+ cervical cancer cell lines (Figure 3C). Together, these data suggest that HR-HPV E6 and E7 regulate MST1 mRNA expression.

**Figure 3.**
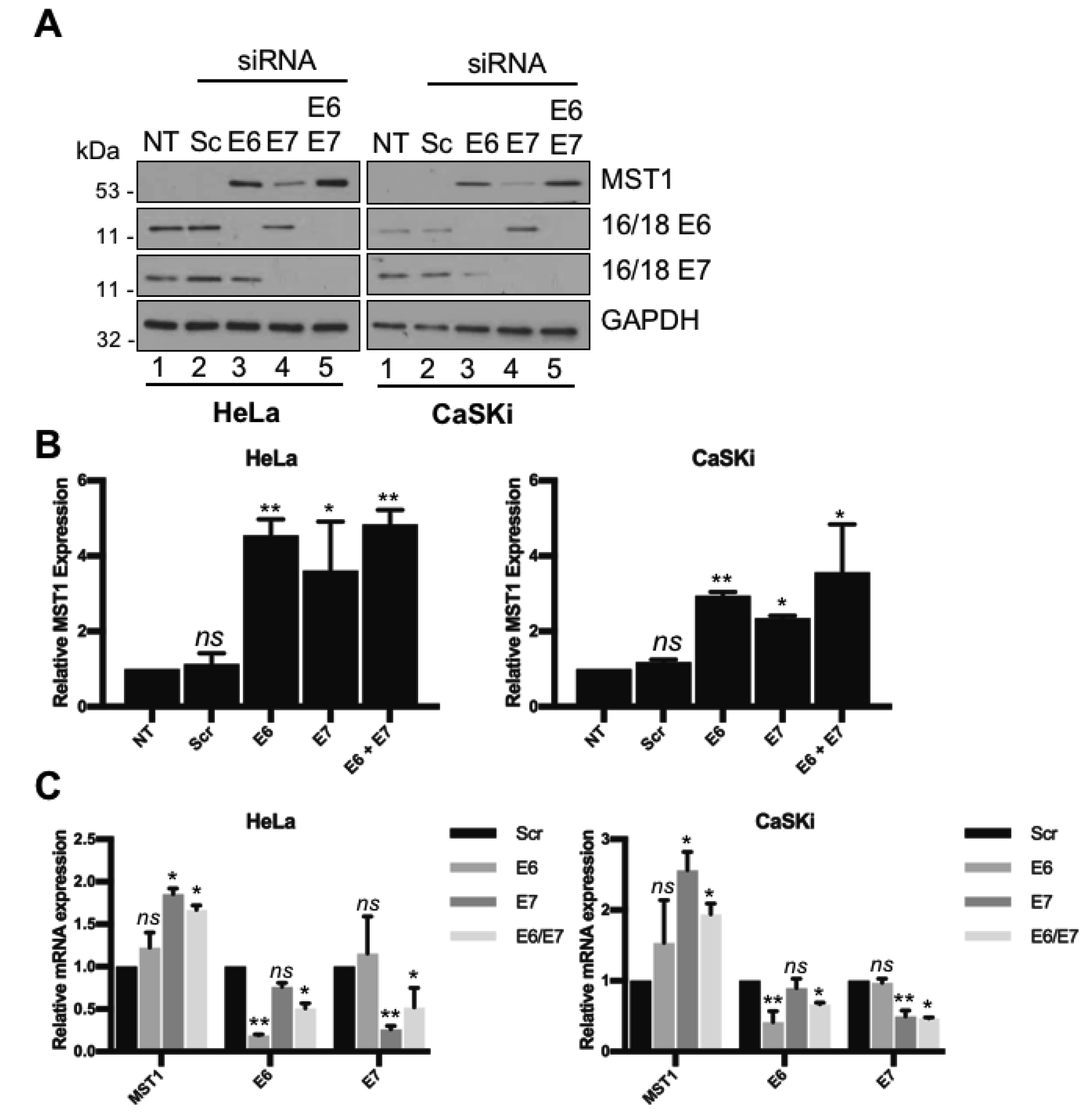
MST1 is regulated at the transcript level in by HR-HPV E6 and E7 oncoproteins. **A)** Representative western blots of HeLa and CaSKi cells transfected with HPV16 or HPV18 specific E6 and/or E7 siRNA, respectively, and analysed for MST1 expression. GAPDH was used as a loading control (N=3). **B)** Quantification of **A** from three independent experiments. **C)** qPCR analysis of MST1 transcript levels in HeLa and CaSKi cells transfected with HPV16 or HPV18 specific E6 and/or E7 siRNA, respectively. Error bars represent the mean +/- standard deviation of a minimum of three biological repeats. *P<0.05, **P<0.01, ***P<0.001 (Student’s t-test).

### MST1/2 over expression activates Hippo signalling in HPV+ cancer cells

To assess the functional consequences of reduced MST1 expression in HPV+cervical cancer, we re-introduced MST1 and its paralogue MST2 into HeLa and CaSKi cells. MST2 shares 75% similarity to MST1 and the two proteins are thought to share some functions (28). Transfection of Myc-tagged MST1 or MST2 led to a significant increase in the phosphorylation of MOB1 and YAP1, indicating that re-introduction of MST1 or MST2 activates the Hippo pathway (Figure 4A and Supplementary Figure 1A). Activated Hippo signalling decreases YAP1 localisation to the nucleus, inhibiting YAP1 transcriptional activity. We therefore used immunofluorescence microscopy to visualise YAP1 levels and localisation in HeLa and CaSKi cells after re-introduction of MST1 and MST2. As shown, MST overexpression resulted in a redistribution of YAP1 from the nucleus to the cytosol (indicated by the white arrows), confirming functional activation of the Hippo pathway by MST1 and MST2 (Figure 4B and Supplementary Figure 1B). To investigate whether the redistribution of YAP1 impacted on transcription, we measured the mRNA levels of several YAP1-dependent genes. Expression of amphiregulin (*AREG),* CD133 *(PROM1),* cyr61 (*cyr61)* and cyclin D1 *(CNND1)* were analysed by qRT-PCR (Figure 4C and Supplementary Figure 1C). Expression of either MST1 or MST2 led to a similar decrease in the mRNA levels of these YAP-dependent genes. To confirm these data, we performed an ELISA for amphiregulin levels in the culture medium of cells overexpressing MST1 or MST2. Amphiregulin levels in the culture medium were reduced on average 50% in cells overexpressing MST1 and 60% in cells expressing MST2 (Figure 4D and Supplementary Figure 1D) These data show that overexpression of MST proteins produces a functionally active Hippo pathway culminating in the exclusion of YAP1 from the nucleus and a reduction in YAP1-dependent gene expression in HPV+cervical cancer cells.

**Figure 4.**
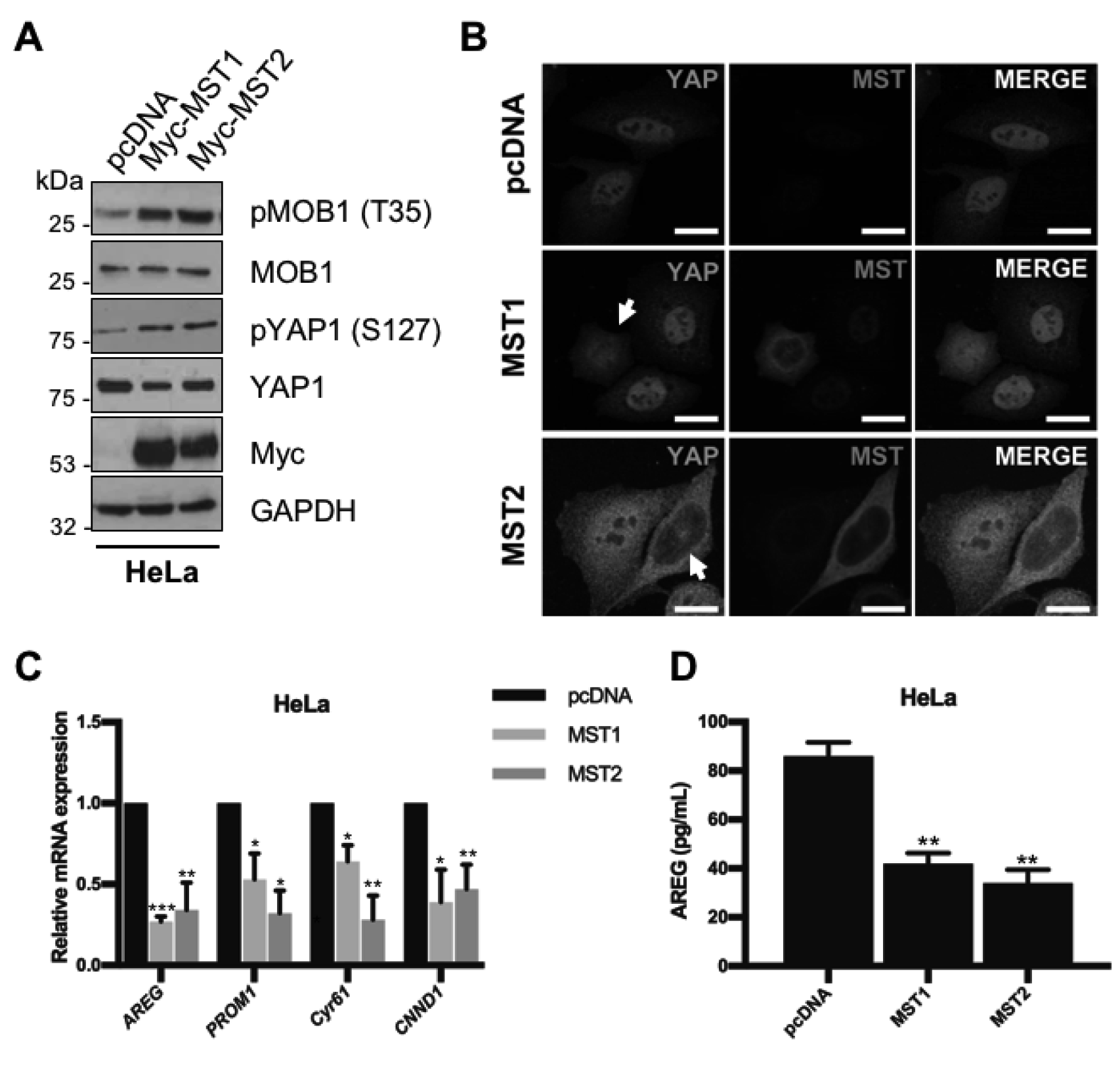
MST1/2 overexpression rescues Hippo signalling in HPV18+ cervical cancer cells. **A)** Representative western blots of MST1/2 overexpression in HeLa cells. Lysates were analysed for the phosphorylation of the MST1/2 substrate MOB1 and the downstream target YAP1. Myc was used to detected successful expression of fusion proteins. GAPDH was used as a loading control (N=3). **B)** Immunofluorescence analysis of MST1/2 overexpression HeLa cells. Cover slips were stained for MST1/2 (red) and YAP1 (green). Nuclei were visualised using DAPI (blue). Images were acquired using identical exposure times. Scale bar, 20 μm. **C)** qPCR analysis of YAP-dependent genes (*AREG, PROM1, Cyr61 and CNND1)* in HeLa cells overexpressing MST1/2. U6 expression was used as a loading control. **D)** ELISA analysis of HeLa cells over expressing MST1 or MST2 for amphiregulin protein in culture medium. Error bars represent the mean +/- standard deviation of a minimum of three biological repeats. *P<0.05, **P<0.01, ***P<0.001 (Student’s t-test).

### MST1/2 inhibits proliferation and cell cycle progression of HPV+ cancer cells

The Hippo pathway negatively regulates tumour progression in a number of cancers. Given this information, we felt it pertinent to investigate the consequences of re-activating Hippo signalling in HPV+ cervical cancer cells by the re-introduction of MST1 or MST2. Compared to HeLa and CaSKi cells transfected with empty expression plasmid, re-introduction of either MST1 or MST2 resulted in a significant decrease in cell proliferation (Figure 5A and Supplementary Figure 2A). Additionally, expression of either MST protein suppressed the ability of HeLa and CaSKi cells to form anchorage-dependant and anchorage-independent colonies (Figure 5B-C and Supplementary Figure 2B-C).

**Figure 5.**
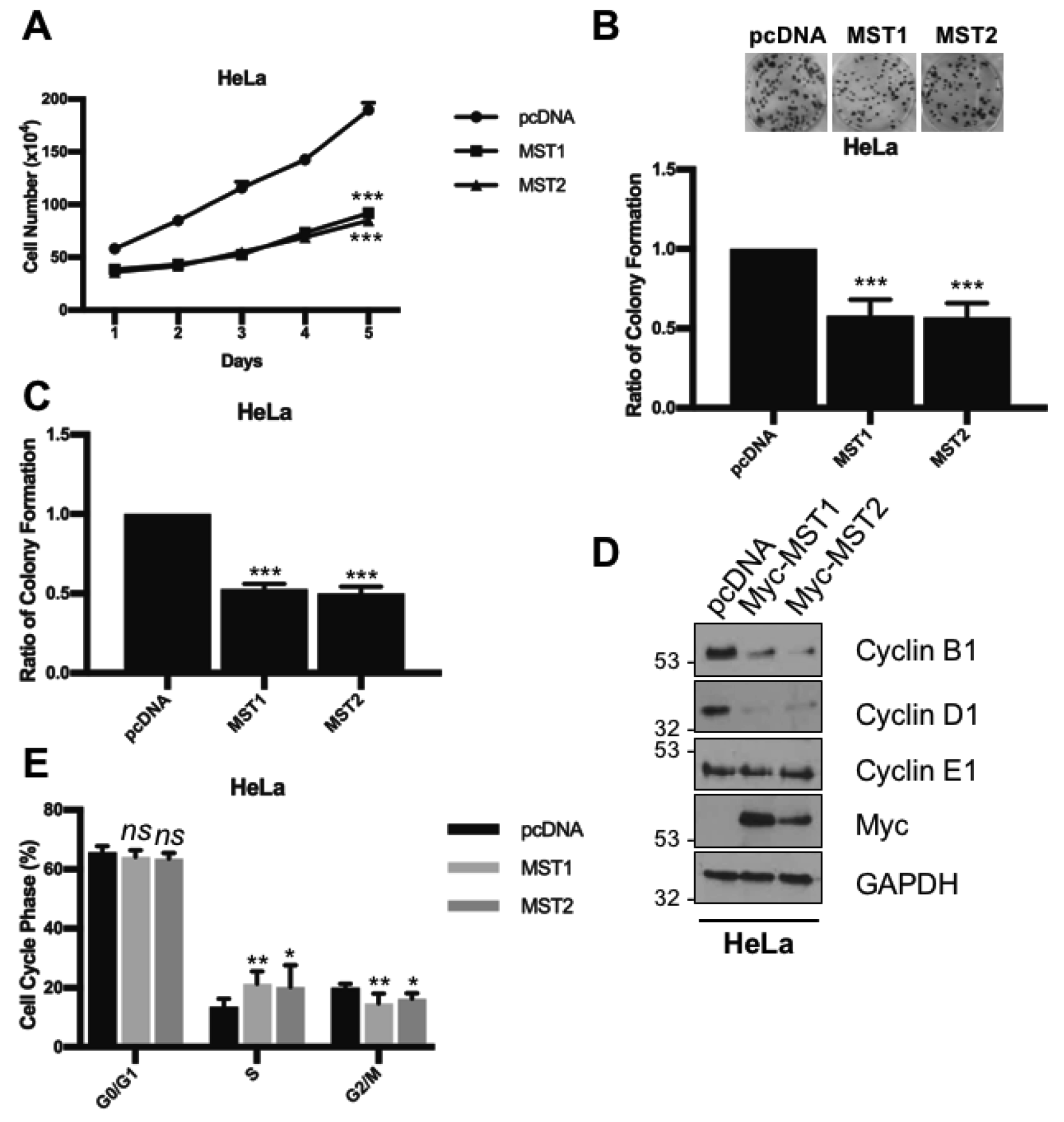
MST1/2 inhibits proliferation and cell cycle progression in HPV+ cervical cancer cells. **A)** Growth curve analysis of HeLa cells overexpressing MST1/2. **B)** Colony formation assay (anchorage dependent growth) of HeLa cells overexpressing MST1/2. **C)** Soft agar assay (anchorage independent growth) of HeLa cells overexpressing MST1/2. **D)** Representative western blots of HeLa cells overexpressing MST1/2 analysed for the expression of cyclin proteins. Myc was used to detected successful expression of fusion proteins. GAPDH was used as a loading control (N=3).. **E)** Flow cytometric analysis of cell cycle profile of HeLa cells overexpressing MST1/2. Error bars represent the mean +/- standard deviation of a minimum of three biological repeats. *P<0.05, **P<0.01, ***P<0.001 (Student’s t-test).

To further evaluate the impact of MST overexpression, we assessed the levels of cyclin proteins, which drive cell proliferation. Upon re-activation of the Hippo pathway, there was a decrease in cyclin B1 and cyclin D1 protein levels but no change in cyclin E expression (Figure 5D and Supplementary Figure 2D). As cyclin B1 and D1 are key markers of cell cycle progression, we investigated the cell cycle status of both HeLa and CaSKi cells overexpressing MST1 and MST2 by flow cytometry (Figure 5E and Supplementary Figure 2E). Analysis revealed an accumulation of cells in S-phase (HeLa MST1 p= 0.004, MST2 p= 0.05; CaSKi MST1 p= 0.03, MST2 p= 0.02) with a corresponding decrease in the number of cells in the G2/M phases, indicating that cells are arrested in S phase.

Together, these data show that Hippo re-activation results in a defect in cell proliferation and the ability of HeLa and CaSKi cells to form colonies in an anchorage dependent or independent manner, highlighting a potential tumour suppressor role for MST proteins in HPV+ cervical cancer cells.

### MST1/2 overexpression does not affect the proliferation of HPV-cancer cells

As we did not observe a significant decrease in MST1 expression in the HPV-C33A cells compared with NHK controls (Figure 1), we assessed whether MST1 or MST2 overexpression could still regulate their proliferation. Despite clear activation of the Hippo pathway in these cells upon MST overexpression (Supplementary Figure 3A-C), we observed a minimal impact upon cell proliferation, colony formation or cell cycle progression (Supplementary Figure 3D-F). This suggests that the tumour suppressor activity of MST proteins is specific to HPV+ cervical cancer cells.

### MST1/2 kinase activity is essential for tumour suppression activity

MST proteins contain a catalytic domain with serine/threonine kinase activity [30]. To investigate whether kinase activity was required for the observed effects of MST overexpression on cell proliferation, we tested the potent MST kinase inhibitor XMU-MP1. Cells overexpressing MST1 or MST2 were treated with XMU-MP1 (1 μM) for 8 housrs prior to analysis. At this dose and time point, no overall effects on cell viability were observed (data not shown). Addition of XMU-MP1 reduced MOB1 and YAP1 phosphorylation to similar levels as seen in the empty plasmid transfected control cells, indicating successful inhibition of MST1 and MST2 kinase activity (Figure 6A and Supplementary Figure 4A). Importantly, treatment of HeLa or CaSKi with XMU-MP1 prevented MST1 or MST2-mediated nuclear exclusion of YAP1 (Figure 6B and Supplementary Figure 4B, white arrows).

**Figure 6.**
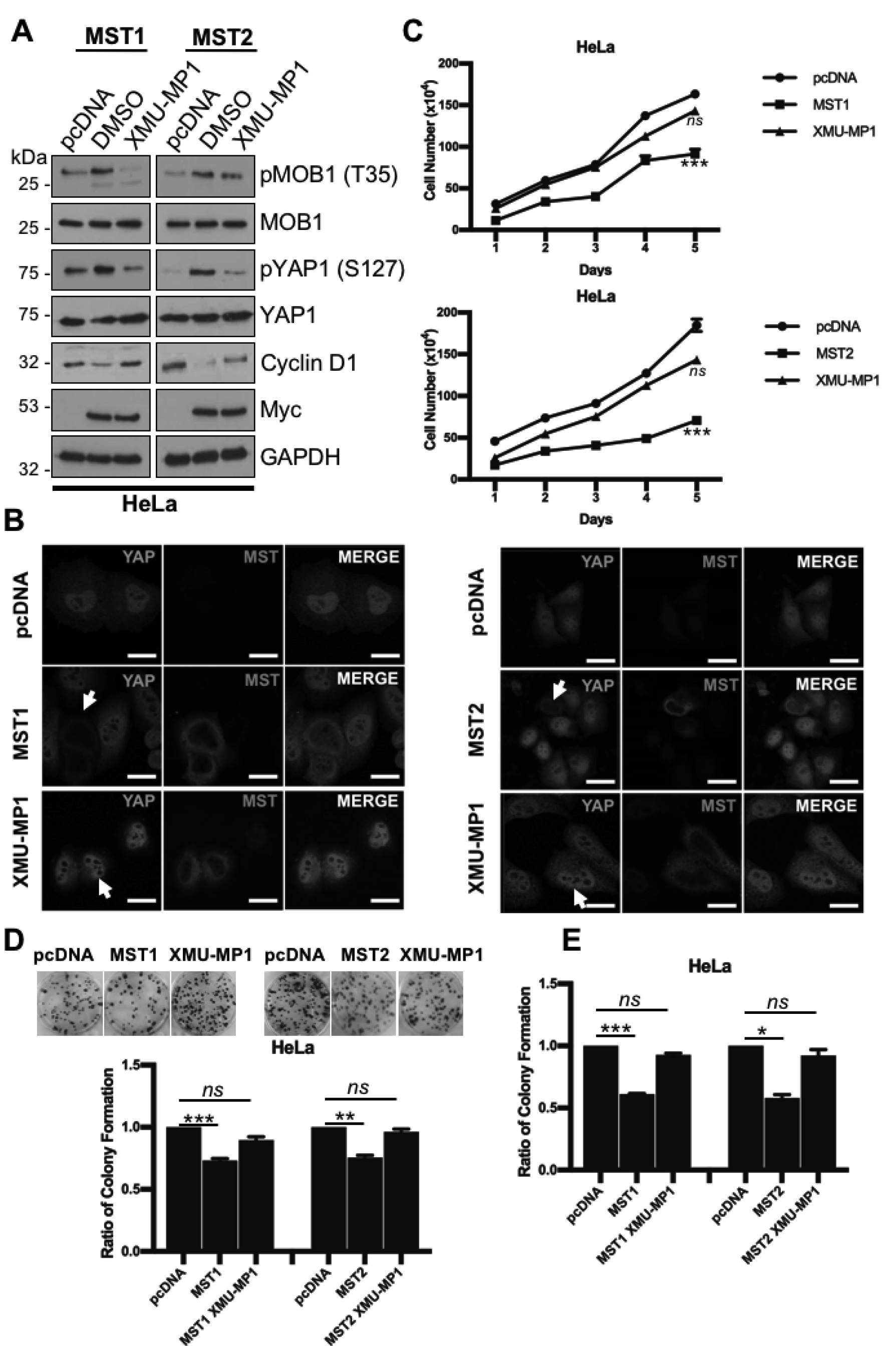
Inhibition of MST1/2 kinase activity prevents the block on proliferation and tumourigenesis. **A)** Representative western blots of MST1/2 overexpression in HeLa cells with or without treatment with XMU-MP1 for 8 hours prior to lysis. Lysates were analysed for the phosphorylation of the MST1/2 substrate MOB1, the downstream target YAP1 and the YAP1 target gene cyclin D1. GAPDH was used as a loading control (N=3). **B)** Immunofluorescence analysis of MST1/2 overexpression in HeLa cells with or without treatment with XMU-MP1 for 8 hours prior to analysis. Cover slips were stained for MST1/2 (red) and YAP1 (green). Nuclei were visualised using DAPI (blue). Images were acquired using identical exposure times. Scale bar, 20 μm. **C)** Growth curve analysis of HeLa cells overexpressing MST1/2 with or without treatment with XMU-MP1 for 8 hours prior to re-seeding. **D)** Colony formation assay (anchorage dependent growth) of HeLa cells overexpressing MST1/2 with or without treatment with XMU-MP1 for 8 hours prior to re-seeding. **E)** Soft agar assay (anchorage independent growth) of HeLa cells overexpressing MST1/2 with or without treatment with XMU-MP1 for 8 hours prior to re-seeding. Error bars represent the mean +/- standard deviation of a minimum of three biological repeats. *P<0.05, **P<0.01, ***P<0.001 (Student’s t-test).

To confirm our observations with XMU-MP1, we introduced either kinase dead MST1 (K59R) or MST2 (K56R) into HPV+ cancer cells. Mutant proteins failed to increase the phosphorylation of MOB compared with wild type (WT) controls (Figure 7A and Supplementary Figure 5A). Furthermore, the mutants failed to redistribute YAP from the nucleus to the cytoplasm (Figure 7B and Supplementary Figure 5B, white arrows), confirming that Hippo pathway activation requires MST1 and MST2 kinase activity in HPV+ cervical cancer cells.

**Figure 7.**
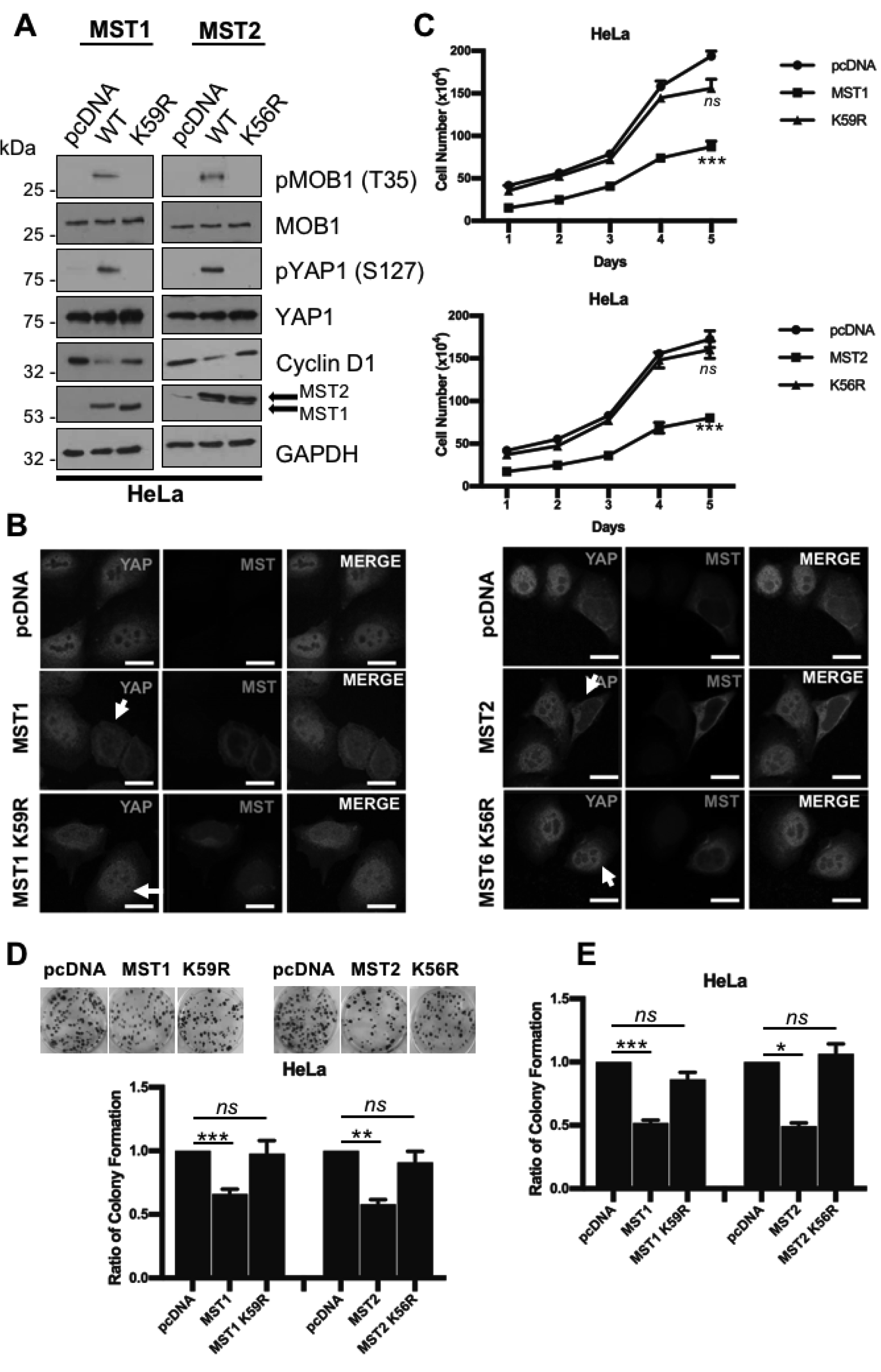
Kinase dead MST1/2 mutants cannot inhibit proliferation and tumourigenesis. **A)** Representative western blots of MST1/2 or KD MST1/2 overexpression in HeLa cells. Lysates were analysed for the phosphorylation of the MST1/2 substrate MOB1, the downstream target YAP1 and the YAP1 target gene cyclin D1. Antibodies against MST1/2 were used to detect successful expression of fusion proteins. GAPDH was used as a loading control (N=3). **B)** Immunofluorescence analysis of MST1/2 or KD MST1/2 overexpression in HeLa cells. Cover slips were stained for MST1/2 (red) and YAP1 (green). Nuclei were visualised using DAPI (blue). Images were acquired using identical exposure times. Scale bar, 20 μm. **C)** Growth curve analysis of HeLa cells overexpressing MST1/2 or KD MST1/2. **D)** Colony formation assay (anchorage dependent growth) of HeLa cells overexpressing MST1/2 or KD MST1/2. **E)** Soft agar assay (anchorage independent growth) of HeLa cells overexpressing MST1/2 or KD MST1/2. Error bars represent the mean +/- standard deviation of a minimum of three biological repeats. *P<0.05, **P<0.01, ***P<0.001 (Student’s t-test).

To further assess the importance of MST kinase activity on cervical cancer cell proliferation, cells overexpressing MST1 or MST2 were incubated with XMU-MP1. XMU-MP1 treatment abolished the effects of MST proteins on cell proliferation (Figure 6C and Supplementary Figure 4C) and colony formation (Figure 6D-E and Supplementary Figure 4D-E). Importantly, similar results were observed when the kinase dead MST1 mutants where overexpressed in HPV+ cells (Figure 7C-E and Supplementary Figure 5C-E). These data strongly suggest that the tumour suppressive functions observed in HPV+ cancer cells require MST kinase activity.

### MST1/2 inhibit the migration and invasion of HPV+ cervical cancer cells

Activated Hippo negatively regulates cancer cell migration and invasion (29,30). To investigate if re-activation of Hippo via the overexpression of MST proteins affects the migration and invasive properties of HPV+ cervical cancer cells, we performed a Transwell^®^ migration assay. MST1 and MST2 overexpression decreased the migration and invasion of both HeLa (Figure 8A and C) and CaSKi cells (Figure 8B and D). To test if this effect was due to MST1/2 kinase activity, we utilised XMU-MP1 and the MST1/2 kinase dead mutants. In contrast to untreated cells overexpressing MST1/2, cells treated with XMU-MP1 or expressing kinase dead MST1/2 mutants did not show a decrease in migration or invasion. Taken together, this demonstrates that MST1/2 negatively regulate migration and invasion of cervical cancer cells in a kinase-dependent manner.

**Figure 8.**
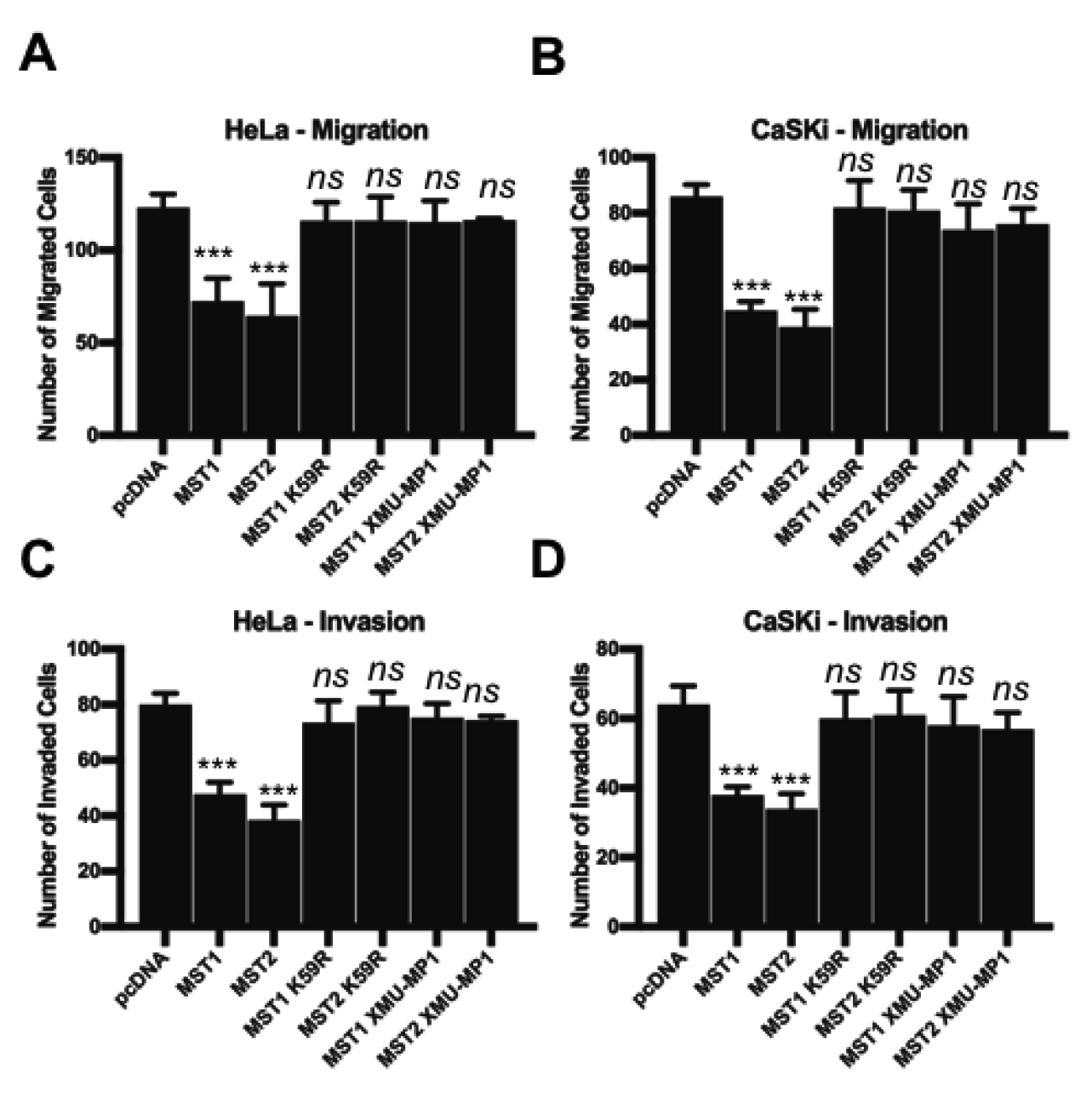
MST1/2 inhibits the migration and invasion of HPV+ cervical cancer cells. **A)-B)** Quantification of the migration of HeLa (**A**) and CaSKi (**B**) cells over expressing MST1/2, KD MST1/2 or MST1/2 after treatment with XMU-MP1 by Transwell^®^ migration assay. The average number of invaded cells per field was calculated from five representative fields per experiment. **C)-D)** Quantification of the invasion of HeLa (**C**) and CaSKi (**D**) cells over expressing MST1/2, KD MST1/2 or MST1/2 after treatment with XMU-MP1 by Transwell^®^ invasion assay. The average number of invaded cells per field was calculated from five representative fields per experiment. Error bars represent the mean +/- standard deviation of a minimum of three biological repeats. *P<0.05, **P<0.01, ***P<0.001 (Student’s t-test).

## Discussion

Despite the current availability of prophylactic vaccines against HPV, there are no specific anti-viral therapies for HPV-associated cancers. Therefore, research is still required to understand the virus-host interactions found in these cancers and to identify the host factors essential for transformation in order to identify targets for therapeutic intervention. The Hippo pathway has emerged as a key signalling pathway deregulated in many cancers (31). As the canonical activator of the Hippo pathway, the MST1 kinase is down-regulated in several cancers and has been shown to serve as a negative regulator of tumourigenesis (16,17). In this study, we used a combination of HPV+ cancer cell lines and patient samples to provide the first evidence that MST1 expression is down regulated in HPV+ cervical cancer.

Our data demonstrates that the HPV E6 and E7 oncoproteins regulate MST1 expression. These oncoproteins have well defined roles to inhibit the function of critical tumour suppressors. The inactivation of p53 and pRb by E6 and E7, respectively, is thought to be a critical step in cervical cancer progression (25,26). In spite of this dogma, E6 mutants exist that are unable to degrade p53 but still transform cells (32). Moreover, inhibition of pRb and its family members is insufficient to initiate cervical cancer (33). These findings highlight the complex nature of virus-mediated transformation and suggest that perturbation of multiple host factors is required for cervical cancer progression.

The Hippo pathway converges on the transcription factor YAP1 - a key mediator of cell proliferation that is deregulated in many cancers, including cervical cancer(19). Importantly, we show that overexpression of MST1 in HPV+ cervical cancer cells leads to an inhibition of YAP1 function, as seen by increased pYAP (S127), and a corresponding decrease in cell proliferation (Figure 4). Recently, it has been demonstrated that HPV16 E6 stabilises the YAP1 protein to enable increased EGFR signalling. This might be achieved by up-regulating expression of EGFR and its ligand amphiregulin, resulting in increased cell proliferation (19). We show that the overexpression of MST1 also down regulates the expression of amphiregulin and this may be a mechanism by which MST1 inhibits cell proliferation.

Our data demonstrates that HPV regulates MST1 mRNA levels (Figure 2 and 3). These findings are supported by clinical data, showing that MST1 mRNA expression negatively correlates with cervical disease progression and data from publicly available databases shows that MST1 mRNA expression is also down regulated in cervical cancer (Figure 1). Thus far the mechanisms by which HPV impairs MST1 expression are not understood. Whilst our knockdown studies show that loss of either E6 or E7 correlates with increased MST1 protein expression, qRT-PCR analysis indicates that E7 is predominantly responsible for repressing MST1 transcription. In head and neck carcinoma expression from the *MST1* gene is down regulated via methylation of its promoter (34). DNA methylation leads to the formation of heterochromatin, an inactive form of DNA, preventing gene expression (35). DNA methylation is an important process in development and alterations in DNA methylation patterns are associated with the development of cancer (36). Given that the HPV oncoproteins have previously been reported to alter levels of host DNA methylation to promote transformation (37), future research will investigate if the *MST1* promoter is methylated in HPV+ cervical cancer to ascertain a mechanism for MST1 mRNA down regulation. Studies will also need to address whether E6 also regulates MST1 protein expression directly. Since E6 interacts with proteins from the ubiquitination system its expression could lead to the degradation of MST1 in cervical cancers.

In summary, our study has demonstrated that the tumour suppressor MST1 is down regulated at the transcript level in HPV positive cervical cancer, leading to dysregulation of the Hippo pathway (Figure 9). We show that overexpression of MST1, and its paralogue MST2, activates the Hippo pathway, resulting in the inhibition of cell proliferation and colony formation ability. Our observations provide further support for targeting the Hippo pathway in HPV positive cervical cancers.

**Figure 9.**
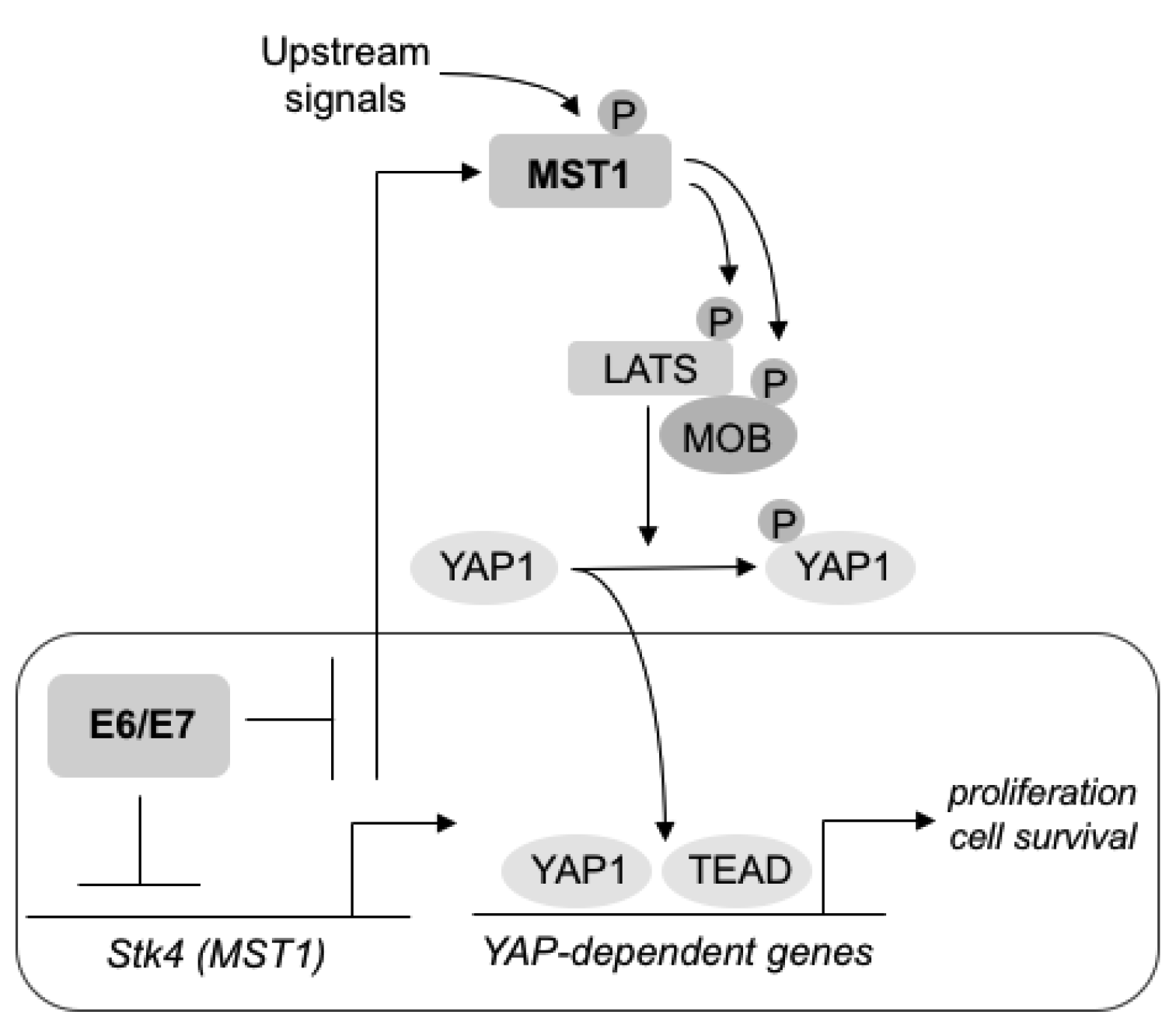
Schematic of HPV-mediated deregulation of the Hippo pathway. Expression of HPV oncoproteins inhibits MST1 mRNA levels, resulting in reduced MST1 protein expression. Loss of MST1 expression correlates with inhibition of the Hippo pathway and activation of the YAP transcription factors, culminating in increased YAP-dependent transcription.

## Acknowledgements

We are grateful to the Wellcome Trust for funding this work via a studentship to E.L.M. (1052221/Z/14/Z). We also acknowledge support from the MRC (MR/K012665/1). We are particularly grateful to Steve Cohen (LMCB, Singapore), Jonathan Chernoff (Fox Chase Cancer Center), Martin Stacey (University of Leeds) and Stephen Griffin (University of Leeds) for generous provision of reagents. We thank the Scottish HPV Investigators Network (SHINE) and Sheila Graham (University of Glasgow) for providing HPV positive cytology and biopsy samples. We would also like to thank Stephen Griffin for helpful comments on the manuscript.

---

**Supplementary Figure 1. MST1/2 overexpression rescues Hippo signalling in HPV16+ cervical cancer cells. A)** Representative western blots of MST1/2 overexpression in CaSKi cells. Lysates were analysed for the phosphorylation of the MST1/2 substrate MOB1 and the downstream target YAP1. Myc was used to detected successful expression of fusion proteins. GAPDH was used as a loading control (N=3).. **B)** Immunofluorescence analysis of MST1/2 overexpression CaSKi cells. Cover slips were stained for MST1/2 (red) and YAP1 (green). Nuclei were visualised using DAPI (blue). Images were acquired using identical exposure times. Scale bar, 20 μm. **C)** qPCR analysis of YAP-dependent genes (*AREG, PROM1, Cyr61 and CNND1)* in CaSKi cells overexpressing MST1/2. U6 expression was used as a loading control. **D)** ELISA analysis of CaSKi cells over expressing MST1 or MST2 for amphiregulin protein in culture medium. Error bars represent the mean +/- standard deviation of a minimum of three biological repeats. *P<0.05, **P<0.01, ***P<0.001 (Student’s t-test).

**Supplementary Figure 2. MST1/2 inhibits proliferation and cell cycle progression in HPV16+ cervical cancer cells. A)** Growth curve analysis of CaSKi cells overexpressing MST1/2. **B)** Colony formation assay (anchorage dependent growth) of CaSKi cells overexpressing MST1/2. **C)** Soft agar assay (anchorage independent growth) of CaSKi cells overexpressing MST1/2. **D)** Representative western blots of CaSKi cells overexpressing MST1/2 analysed for the expression of cyclin proteins. Myc was used to detected successful expression of fusion proteins. GAPDH was used as a loading control (N=3). **E)** Flow cytometric analysis of cell cycle profile of CaSKi cells overexpressing MST1/2. Error bars represent the mean +/- standard deviation of a minimum of three biological repeats. *P<0.05, **P<0.01, ***P<0.001 (Student’s t-test).

**Supplementary Figure 3. MST1/2 does not inhibit proliferation and cell cycle progression in HPV-cervical cancer cells. A)** Representative western blots of MST1/2 overexpression in C33A cells. Lysates were analysed for the phosphorylation of the MST1/2 substrate MOB1 and the downstream target YAP1. Myc was used to detected successful expression of fusion proteins. GAPDH was used as a loading control (N=3). **B)** Immunofluorescence analysis of MST1/2 overexpression in C33A cells. Cover slips were stained for MST1/2 (red) and YAP1 (green). Nuclei were visualised using DAPI (blue). Images were acquired using identical exposure times. Scale bar, 20 μm. **C)** qPCR analysis of YAP-dependent genes (*AREG, PROM1, Cyr61 and CNND1)* in C33A cells overexpressing MST1/2. U6 expression was used as a loading control. **D)** Growth curve analysis of C33A cells overexpressing MST1/2. **E)** Colony formation assay (anchorage dependent growth) of C33A cells overexpressing MST1/2. **F)** Flow cytometric analysis of cell cycle profile of C33A cells overexpressing MST1/2. Error bars represent the mean +/- standard deviation of a minimum of three biological repeats. *P<0.05, **P<0.01, ***P<0.001 (Student’s t-test).

**Supplementary Figure 4. Inhibition of MST1/2 kinase activity prevents the block on proliferation and tumourigenesis A) A)** Representative western blots of MST1/2 overexpression in CaSKi cells with or without treatment with XMU-MP1 for 8 hours prior to lysis. Lysates were analysed for the phosphorylation of the MST1/2 substrate MOB1, the downstream target YAP1 and the YAP1 target gene cyclin D1.GAPDH was used as a loading control (N=3). **B)** Immunofluorescence analysis of MST1/2 overexpression in CaSKi cells with or without treatment with XMU-MP1 for 8 hours prior to analysis. Cover slips were stained for MST1/2 (red) and YAP1 (green). Nuclei were visualised using DAPI (blue). Images were acquired using identical exposure times. Scale bar, 20 μm. **C)** Growth curve analysis of CaSKi cells overexpressing MST1/2 with or without treatment with XMU-MP1 for 8 hours prior to re-seeding. **D)** Colony formation assay (anchorage dependent growth) of CaSKi cells overexpressing MST1/2 with or without treatment with XMU-MP1 for 8 hours prior to re-seeding. **E)** Soft agar assay (anchorage independent growth) of CaSKi cells overexpressing MST1/2 with or without treatment with XMU-MP1 for 8 hours prior to re-seeding. Error bars represent the mean +/- standard deviation of a minimum of three biological repeats. *P<0.05, **P<0.01, ***P<0.001 (Student’s t-test).

**Supplementary Figure 5. Kinase dead MST1/2 mutants cannot inhibit proliferation and tumourigenesis A) A)** Representative western blots of MST1/2 or KD MST1/2 overexpression in CaSKi cells. Lysates were analysed for the phosphorylation of the MST1/2 substrate MOB1, the downstream target YAP1 and the YAP1 target gene cylin D1. Antibodies against MST1/2 were used to detect successful expression of fusion proteins. GAPDH was used as a loading control (N=3). **B)** Immunofluorescence analysis of MST1/2 or KD MST1/2 overexpression in CaSKi cells. Cover slips were stained for MST1/2 (red) and YAP1 (green). Nuclei were visualised using DAPI (blue). Images were acquired using identical exposure times. Scale bar, 20 μm. **C)** Growth curve analysis of CaSKi cells overexpressing MST1/2 or KD MST1/2. **D)** Colony formation assay (anchorage dependent growth) of CaSKi cells overexpressing MST1/2 or KD MST1/2. **E)** Soft agar assay (anchorage independent growth) of CaSKi cells overexpressing MST1/2 or KD MST1/2. Error bars represent the mean +/- standard deviation of a minimum of three biological repeats. *P<0.05, **P<0.01, ***P<0.001 (Student’s t-test).

